# DeepDigest: prediction of protein proteolytic digestion with deep learning

**DOI:** 10.1101/2020.03.13.990200

**Authors:** Jinghan Yang, Zhiqiang Gao, Xiuhan Ren, Jie Sheng, Ping Xu, Cheng Chang, Yan Fu

**Affiliations:** National Center for Mathematics and Interdisciplinary Sciences, Key Laboratory of Random Complex Structures and Data Science, Academy of Mathematics and Systems Science, Chinese Academy of Sciences, Beijing 100190, China; State Key Laboratory of Proteomics, Beijing Proteome Research Center, National Center for Protein Sciences (Beijing), Beijing Institute of Lifeomics, Beijing 102206, China; School of Mathematical Sciences, University of Chinese Academy of Sciences, Beijing 100049, China; China University of Mining & Technology, Beijing 100083, China

## Abstract

In shotgun proteomics, it is essential to accurately determine the proteolytic products of each protein in the sample for subsequent identification and quantification, because these proteolytic products are usually taken as the surrogates of their parent proteins in the further data analysis. However, systematical studies about the commonly used proteases in proteomics research are insufficient, and there is a lack of easy-to-use tools to predict the digestibilities of these proteolytic products. Here, we propose a novel sequence-based deep learning model – DeepDigest, which integrates convolutional neural networks and long-short term memory networks for digestibility prediction of peptides. DeepDigest can predict the proteolytic cleavage sites for eight popular proteases including trypsin, ArgC, chymotrypsin, GluC, LysC, AspN, LysN and LysargiNase. Compared with traditional machine learning algorithms, DeepDigest showed superior performance for all the eight proteases on a variety of datasets. Besides, some interesting characteristics of different proteases were revealed and discussed.

## INTRODUCTION

The protein proteolytic digestion is a key step in shotgun proteomics, and the proteolytic products of their parent proteins are usually taken as surrogates in subsequent identification and quantification protocol^1^. Till now, trypsin is the most popular and widely used protease in proteomics because of its high specificity, ease of use and cost efficiency, and it generates mass spectrometry (MS)-favored proteolytic peptides which have detectable masses and charges by tandem mass spectrometry (MS/MS)^2,3^. However, the sole use of trypsin may hinder our ability to explore the full proteome, missing particular cleavage sites or protein segments^2^. Therefore, some alternative proteases are also used to complement trypsin, such as ArgC, chymotrypsin, GluC, LysC, AspN, LysN and LysargiNase. But, regardless of which protease is used, the proteolytic digestion can be hardly complete. For example, it was reported that about 40% of unique identified peptides digested by trypsin contained one or more missed cleavages^4^.

In MS/MS-based protein identification by database searching, it is necessary to match the experimental spectrum with the theoretical spectra of the in silico digested peptides. Thus, the ultimate goal of in silico digestion is to simulate the experimental proteolytic process as accurately as possible^5^. In the label-free protein quantification, the abundance of a protein is usually inferred from the mass spectrometric signals of its digested peptides. Thus, the quantitative value of a protein would be underestimated if incomplete proteolytic digestion is not considered. Therefore, accurate prediction of proteolytic digestion is essential to both identification and quantification of proteins.

By now, a series of studies on predicting the protein cleavage sites have been published, especially for the trypsin protease^6,7^. For example, Keil et al.^7,8^ summarized the rules, called “Keil rules”, of missed cleavage sites for trypsin, mainly consisting of 1) Arginine (R)/Lysine (K) is followed by Proline (P); 2) two successive R/K is close to each other; 3) Aspartic acid/glutamic acid residues is adjacent to the positively charged residue. However, the “Keil rules” were empirical and not based on MS data. Rodriguez et al.^9^ reported discrepant results about these rules: the number of peptides produced by supposedly “illegitimate” [RK].[P] cleavages was higher than the number of peptides produced by legitimate [RK].[C] cleavages. Actually, it should be noted that proteolytic digestion is a complicated probability problem, i.e., one digestion site may be cleaved with a certain probability, more than a simple binary “cut or not cut” problem. Unfortunately, these empirical rules cannot accurately calculate the peptide digestibility, which is defined as the probability of the peptide being digested by the protein digestion process^10^. Thus, computational models and algorithms were introduced. Siepen et al.^11^ predicted missed cleavage sites by information theory, with up to 90% accuracy using amino acid sequences alone. Lawless et al.^12^ then built a missed cleavage site predictor using the support vector machine (SVM) algorithm with an overall validated area under the ROC curve (AUC) of 0.88. CP-DT was developed to predict tryptic cleavages based on decision tree ensembles and achieved an AUC of at least 83% on all the test datasets^7^. Despite these developments, the prediction accuracy of tryptic sites is still far from satisfactory; moreover, there are fewer related works for other proteases than trypsin^3,13^. Meanwhile, there is not yet a specialized tool for predicting cleavage sites of various common proteases in proteomics.

In this study, we present a deep learning-based protein proteolytic digestion predictor, named DeepDigest, which integrates the merits of convolutional neural networks (CNNs) and long-short term memory (LSTM) networks. Two convolutional layers extract the local sequence features and an LSTM layer explores the long-term dependencies between amino acids. On a variety of test sets, DeepDigest showed superior performance over traditional machine learning algorithms (logistic regression, random forest and SVM) with the same training sets. Our experiments involved eight proteases, i.e., trypsin, ArgC, chymotrypsin, GluC, LysC, AspN, LysN and LysargiNase^14–21^. The results demonstrated that DeepDigest retained good predictive power on independent test sets. In addition, some interesting characteristics of different proteases were revealed and discussed. For example, our results confirmed that the C-terminal amino acids of the cleavage sites had greater impact on proteolytic digestion than N-terminal amino acids for the C-terminal proteases, and the opposite for the N-terminal proteases.

## EXPERIMENTAL SECTION

### Datasets and Database Searching

Nineteen public large-scale datasets, covering eight typical proteases, i.e., trypsin, ArgC, chymotrypsin, GluC, LysC, AspN, LysN, and LysargiNase, were utilized in this study (Table 1). These datasets were from samples of four organisms including *E. coli*, yeast, mouse and human. The proteolytic digestion was performed overnight as described in their original papers^14,16–21^. To learn adequate sequence patterns, the first eight datasets were used as training sets, while the other eleven were test sets. The raw MS files were downloaded and reanalyzed as follows.

**Table 1.**
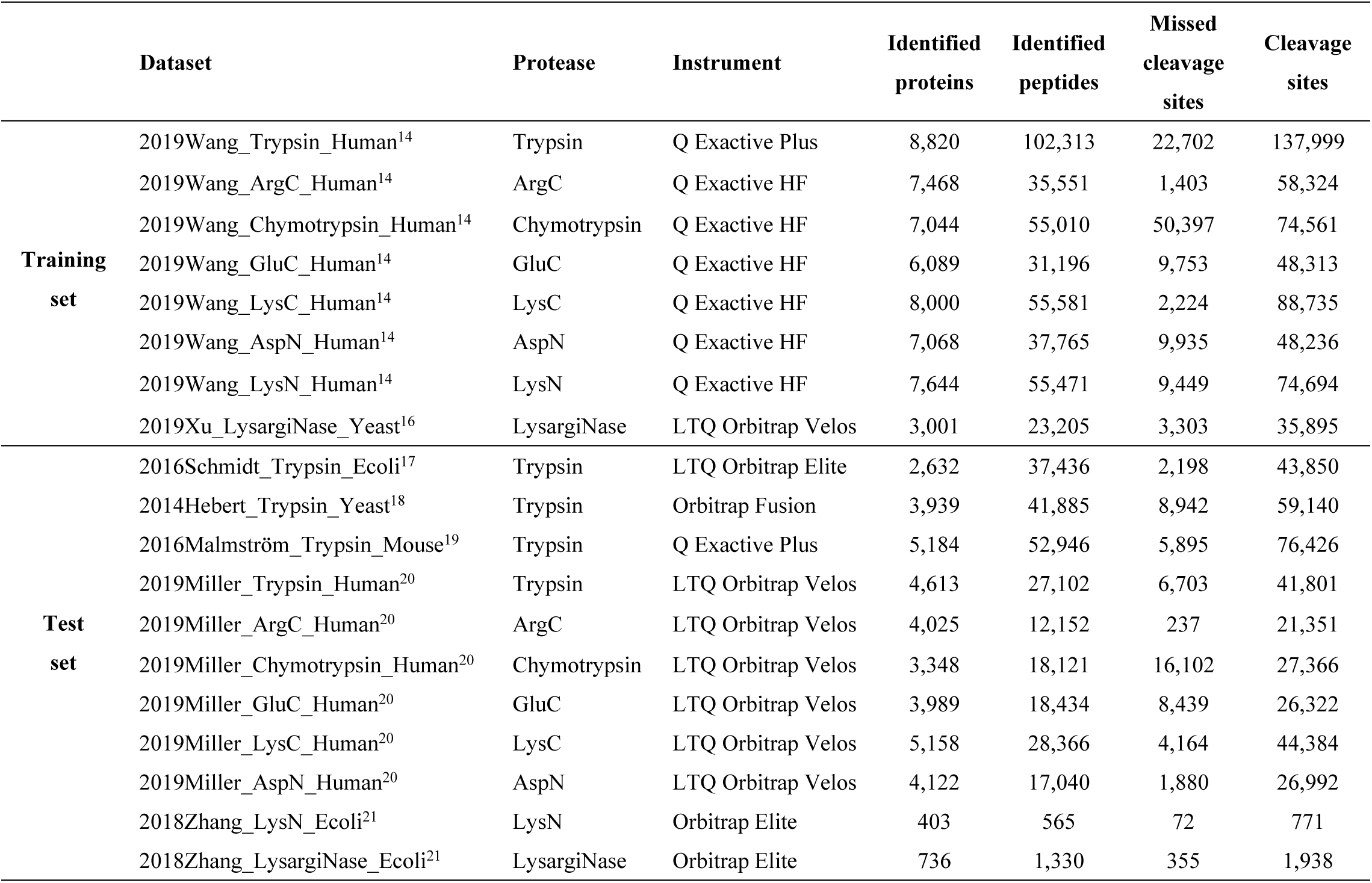
Summary of the public datasets used for training and testing DeepDigest.

Mass spectra in each dataset were searched against the corresponding organism sequences from the UniProt database using the Andromeda search engine^22^ in the software MaxQuant^23^ (version 1.6.0.1). For the two *E. coli* datasets in Zhang et al.’s paper^21^, propionamidation on cysteine, ethanolamine protection on aspartate and glutamate, dimethylation on peptide N-term, and dimethylation on lysine were specified as fixed modifications; oxidation on methionine, dimethylation on protein N-term, and ethanolamine protection on protein C-term were specified as variable modifications; the precursor mass tolerance was set to 20 ppm for the first search (for the identification of the maximum number of peptides for mass and retention time calibration) and 10 ppm for the main search (for the refinement of the identifications). For the other datasets, carbamidomethylation on cysteine was set as a fixed modification; oxidation on methionine and protein N-terminal acetylation were set as variable modifications; the precursor mass tolerance was set to 20 ppm for the first search and 4.5 ppm for the main search. Protein sequences were theoretically digested by one of the eight proteases with up to two missed cleavages allowed, and the false discovery rate was set to 1% at both peptide-spectrum match and protein levels.

### Data Preprocessing

Identified peptides were first mapped to their corresponding proteins; then, the digestion information of each cleavage site in the identified proteins was collected, involving the spectral counts (SCs) of the peptides observed on the N-terminal (SC_N_) and C-terminal (SCC) of the cleavage site, and the SC of the peptides containing this cleavage site as a missed cleavage site (SC_M_). Afterwards, the cleavage site was labeled as a positive site if (1) SC_N_ or SCC was at least 1 and (2) SC_M_ was zero, and as a negative site if (1) both SC_N_ and SCC were zero and (2) SC_M_ was at least 1. Previous studies had demonstrated that the digestibility of a cleavage site was mainly influenced by its adjacent amino acids^7,10,12,24^. Therefore, for each positive or negative site, fifteen adjacent residues on both sides were extracted, resulting in a 31-mer including the cleavage site^25^. Moreover, if there were not enough amino acids on the N- or C-terminal of a protein, the character Z was added to make up a 31-mer. Finally, each character in each 31-mer was encoded by alphabetical order (Table S1) for convenience. The coding order had no impact on results.

### Architecture of DeepDigest Model

As shown in Figure 1, DeepDigest integrates CNNs and LSTM networks to detect discriminatory patterns around the cleavage sites and capture the long-term dependencies between amino acids. With the embedding approach, the distributed representation of amino acids was learned. Moreover, in order to overcome the imbalance problem that there were more cleavage sites than missing ones, DeepDigest was trained with a weighted binary cross entropy loss function whose weights were empirically tuned ^26,27^.

**Figure 1.**
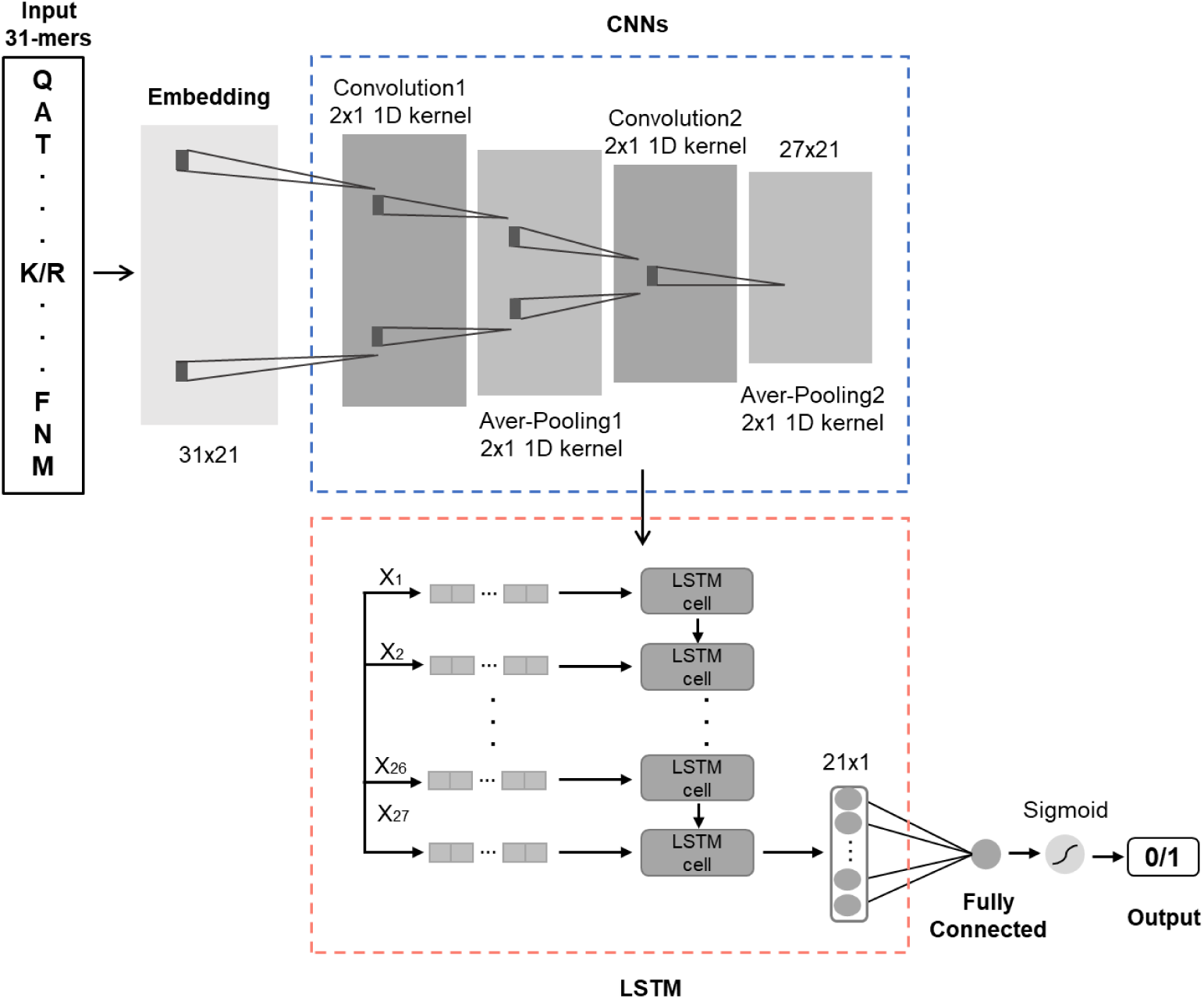
The network architecture of DeepDigest.

Representation learning is one of the critical abilities of deep learning. Actually, word embedding is a set of feature learning techniques, such as word2vec^28,29^. It is commonly used in natural language processing, mapping words from the vocabulary to vectors of real numbers. Sparse representations, e.g. one-hot representation, may cause a lot of difficulties while training deep learning models. Thus, it is necessary to use word embedding, which is capable of learning denser continuous feature vectors, also known as distributed representation. In DeepDigest, each encoded character in the 31-mer is projected into a 21-dimensional vector by the embedding layer^30^. In this way, more sensible representation can be learned and the local interaction between amino acids can be deeply explored by the convolution layers.

The embedding outputs are then fed to CNNs with two convolutional layers and two average pooling layers. The convolutional operator is supposed to filter as many local features as possible with several different kernels, which are often called the “feature detector”^31^. Each of the two convolutional layers in our model contains 21 kernels of length 2 with a stride of 1, followed by an activation with Rectified Linear Unit (ReLU)^32^. The average pooling layer whose window size is 2 with a stride of 1 is applied after each convolutional layer. With the output of the CNNs, DeepDigest has fully extracted critical local features with shape 27×21.

Last but not least, the following LSTM networks as a special kind of recurrent neural networks, can capture dependencies across a longer distance among amino acids without gradient vanishing or exploding problems. Particularly, each LSTM cell can be seen as a projection with nonlinear transformations. The output length is set to 21, which is equal to the number of useful characters. The final layer with one neuron is activated by a sigmoid function to predict the probability of proteolytic digestion.

## RESULTS AND DISCUSSION

### Performance of DeepDigest

To evaluate the prediction performance of DeepDigest, the AUCs, F1 scores and Matthews correlation coefficients (MCCs) of 10-fold cross-validations were calculated on the training sets and shown in Table S2. In order to demonstrate the superiority of DeepDigest over traditional machine learning algorithms, we compared it with logistic regression (LR), random forest (RF), and SVM. The AUCs of DeepDigest were 0.968∼0.983 as shown in Figure 2, F1 scores and MCCs were 0.639∼0.904 and 0.631∼0.839 respectively, as shown in Table S2. For each protease, DeepDigest had the highest values of AUCs, F1 scores and MCCs. Especially on the 2019Wang_Chymotrypsin_Human dataset, the AUCs, F1 scores and MCCs of DeepDigest had respectively increased by 9.7%, 19.1%, 37.5% compared to LR, by 10.2%, 24.0%, 42.4% compared to RF, and by 8.7%, 17.7%, 33.6% compared to SVM. These results showed that our model had a superior learning performance on various proteases.

**Figure 2.**
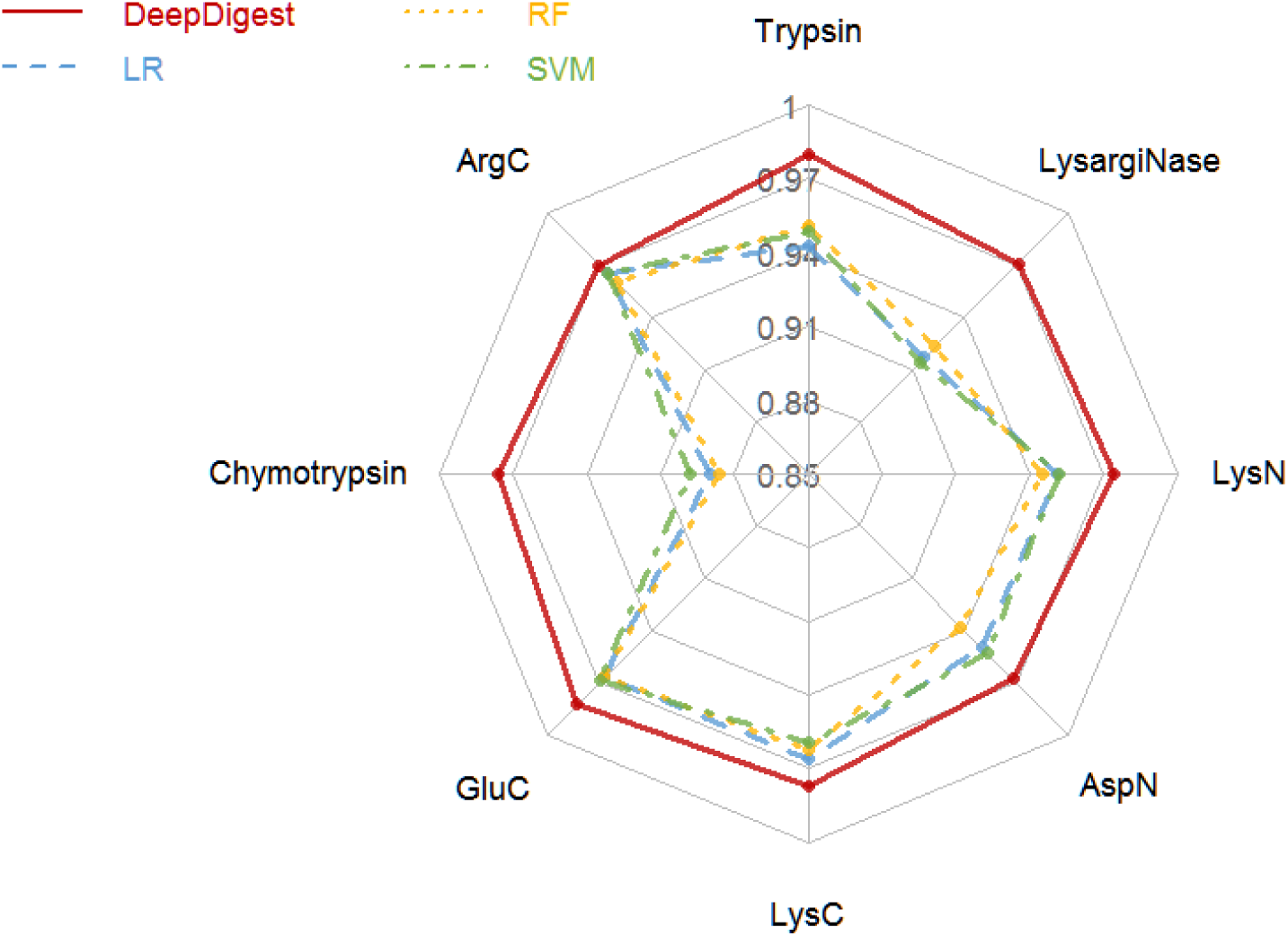
Ten-fold cross-validation AUCs of DeepDigest on the eight training sets, compared with logistic regression (LR), random forest (RF), and support vector machine (SVM).

To evaluate the generalization ability of DeepDigest, the four models (DeepDigest, LR, RF and SVM) trained on the training sets, were tested on the eleven independent test sets as shown in Table 1. The AUCs, F1 scores, and MCCs were shown in Table S3. The AUCs of DeepDigest were between 0.809 and 0.977, while the AUCs of LR, RF and SVM were 0.736∼0.965, 0.779∼0.962, and 0.744∼0.972 respectively as shown in Figure 3. These results demonstrated that DeepDigest had superior generalization ability for various proteases.

**Figure 3.**
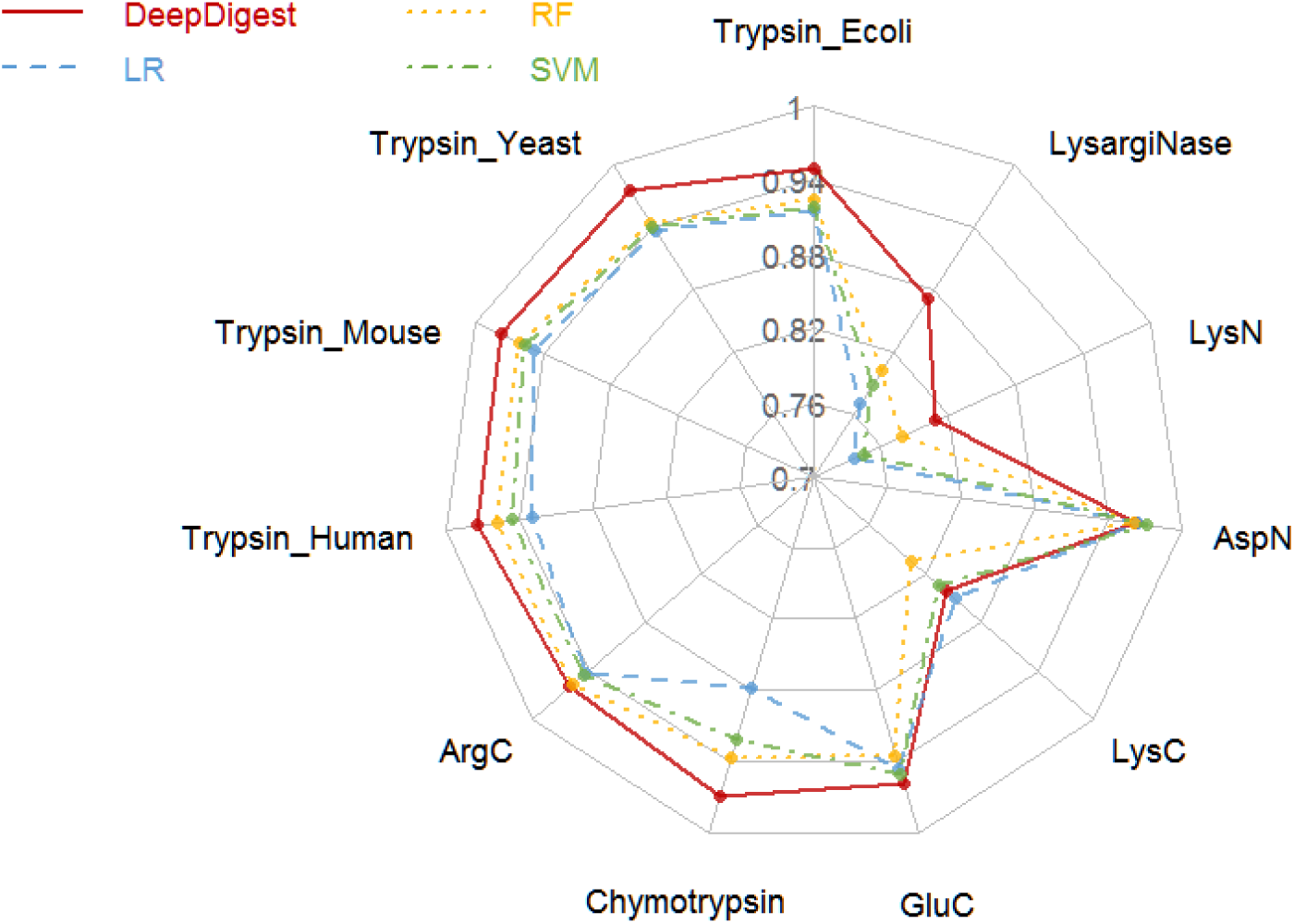
Test AUCs of DeepDigest on the eleven independent test sets, compared with logistic regression (LR), random forest (RF), and support vector machine (SVM).

### Comparison with Existing Tool

So far, the available tools for predicting trypsin cleavage sites or missed cleavage sites are MS-Digest^33^, PeptideCutter^34^, CP-DT^7^ and MC:pred^6^. They are all provided in the form of web-servers. Specifically, MS-Digest and PeptideCutter can only provide possible cleavage sites without peptide digestibility. Moreover, PeptideCutter and CP-DT can only take one protein sequence as input each time, making it impossible for large-scale data analysis. At last, only MC:pred was compared with DeepDigest. It should be noted that MC:pred cannot process amino acids that are represented by uncommonly used alphabet, for example, X, O, etc. Therefore, the proteins containing these amino acids were removed before running MC:pred. The comparison of DeepDigest and MC:pred was carried out on the four independent tryptic test sets from four different organisms (*E. coli*, Yeast, Mouse and Human). Here, DeepDigest was trained on the 2019Wang_Trypsin_Human dataset, and the predicted peptide digestibility was also calculated^10^. As shown in Figure S1, DeepDigest had AUCs of 0.967, 0.988, 0.988 and 0.985 on the four test sets respectively, while MC:pred had AUCs of 0.961, 0.945, 0.972 and 0.935. Overall, DeepDigest had the higher prediction accuracy than MC:pred.

### Length of Input Sequence Fragments

In this study, the hypothesis is that the digestion of the cleavage site is mainly influenced by the surrounding amino acids. Several related algorithms have been developed based on this assumption^6,11,25^. However, it is still unknown how far away the amino acids from the cleavage sites will affect the digestion behavior. Thus, centering on the potential cleavage sites, 1, 2, 3… 15 amino acids on both sides were respectively extracted to form amino acid sequence fragments of length 3, 5, 7… 31, which were taken as the inputs of DeepDigest. This experiment was performed on the training sets. As shown in Figure S2(A), for various proteases, AUCs of DeepDigest gradually increased as the length of input sequence fragments increased; and when the length reached 31, the prediction ability of DeepDigest was basically saturated. Furthermore, to ensure the rationality of the input length, the same experiments were performed on the same datasets with other two traditional machine learning methods – LR and RF, and the similar trend was present in Figure S2(B-C). Specifically, we made a random 80-20% train-test split for each dataset. These results showed that the length of input sequence fragments was independent of models. Therefore, the number of amino acids extracted on both sides of the cleavage sites was fixed to 15 for this study.

### Effect of Amino Acids from Different Sides

Furthermore, to explore the mechanisms of proteolytic digestion of various proteases, we conducted an in-depth analysis of the effect of adjacent amino acids on the cleavage site. Specifically, 1, 2, 3… or 15 amino acids from the left/right side of the cleavage sites were also used to form the input sequence fragments of 2, 3, 4… or 16-mer (cleavage site included). Thus, input sequence fragments with the increasing number of amino acids on the both sides and the left/right side of the cleavage sites were generated. As shown in Figure 5, the AUCs of DeepDigest, fed with input sequence fragments of amino acids on both sides, were the highest on all the training sets for all the eight proteases. Besides, it indicated that the C-terminal amino acids of the cleavage sites had greater impact on the cleavage than N-terminal amino acids for the C-terminal proteases (Figure 4A-E), and the opposite for the N-terminal proteases (Figure 4F-H). Anyway, considering that the mechanisms of proteolytic digestion are related to the structures and physiochemical properties of proteins, the cleavage behavior indeed could be more affected by the followed amino acids of the cleavage sites.

**Figure 4.**
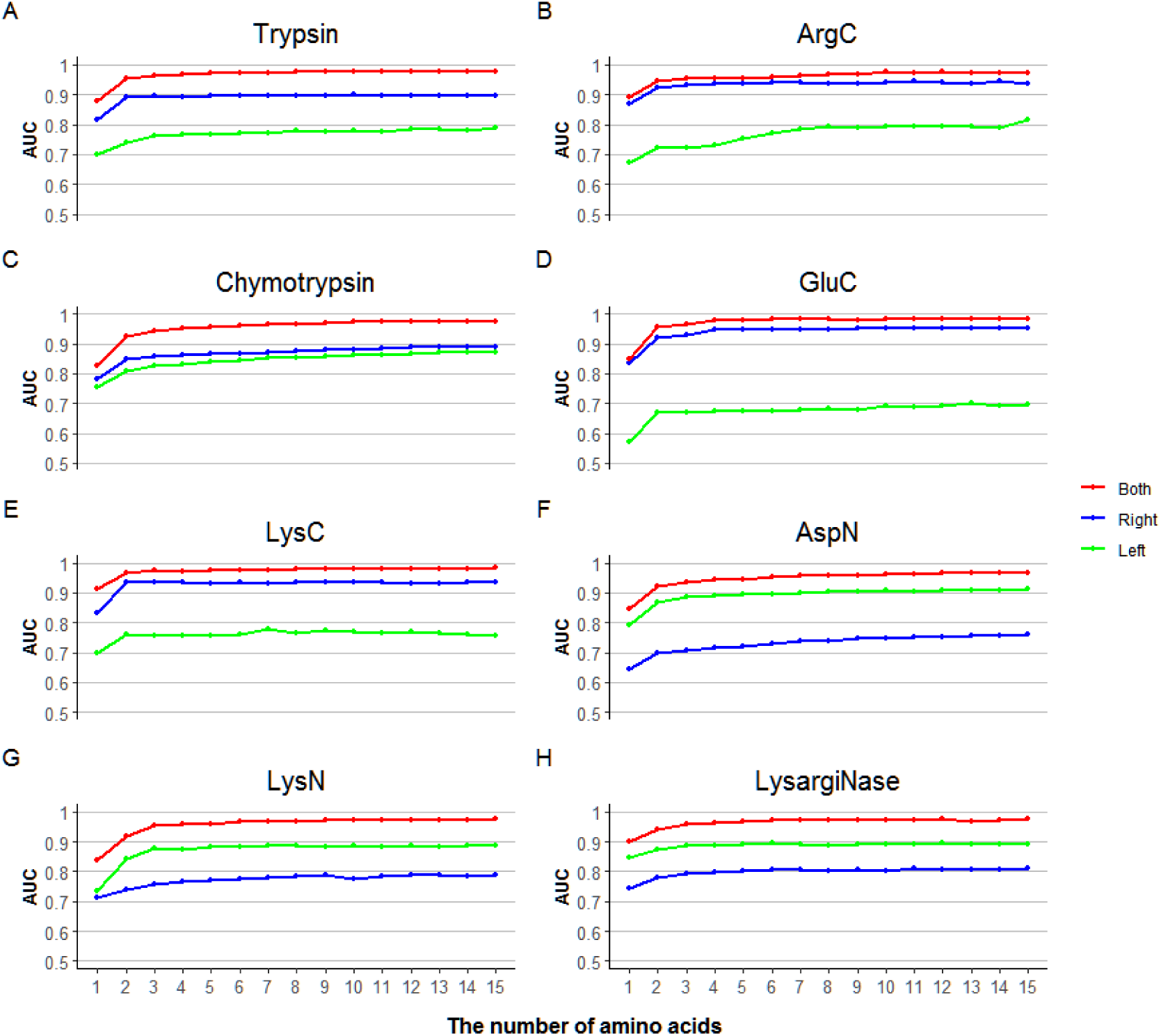
The AUCs of DeepDigest with the increasing number of amino acids on the Both sides and from the Right/Left side of the cleavage site with eight different proteases (A) Trypsin, (B) ArgC, (C) Chymotrypsin, (D) GluC, (E) LysC, (F) AspN, (G) LysN and (H) LysargiNase.

### Sequence Motif Analysis

To explore the features surrounding a cleavage site or a missed cleavage site, the sequence logo diagrams of all the input 31-mers were displayed with the pLogo^35^. As shown in Figure S3, for the proteases such as Trypsin, ArgC, LysC, LysN and LysargiNase, whose cleavage sites are basic amino acids (lysine or arginine), the cleavage sites tended to be missed if glutamic acid and aspartic acid were significantly overrepresented. Moreover, it was likely to be a missed cleavage site if there was proline at the position -2, -1, +1, or +2 of the site, specifically for Trypsin (position +1, +2), ArgC (position +1), GluC (position -2, +1, +2), LysC (position +1, +2), AspN (position -1, +1) and LysN (position -2, +1). Giansanti et al.^2^ showed that if acidic amino acids (glutamic acid and aspartic acid) were at the position -1, +1, +2 of a cleavage site, this site was likely to be missed for ArgC. Our results of ArgC further showed that there were glutamic acid and aspartic acid simultaneously at the position +3, +4 of a missed cleavage site.

## CONCLUSIONS

In this study, we developed a novel model based on deep learning, named DeepDigest, to predict proteolytic cleavage sites and the peptide digestibility for eight commonly used proteases. To our knowledge, it is the first time that the deep learning technique is used to solve the digestibility prediction problem, and it turned out that the deep learning model worked better than the traditional machine learning methods. Furthermore, there are a lot of important applications of peptide digestibility in experimental proteomics. For example, it has been utilized to improve peptide mass fingerprinting scoring^11,36^ or peptide detectability prediction^10^. This study provided an easy-to-use tool, DeepDigest, to predict the digestibilities of peptides digested by multiple proteases, which could have contribution to the selection of surrogate peptides, and protein identification and quantitation. Because peptides with higher predicted digestibility tend to be selected as surrogates of their parent proteins.

## ACKNOWLEDGEMENT

This study was supported by the NCMIS CAS and the State Key Laboratory of Proteomics [SKLP-Y201802].

