## Supporting Information for "DeepDigest: prediction of protein proteolytic digestion with deep learning"

to

### Supplementary Figures

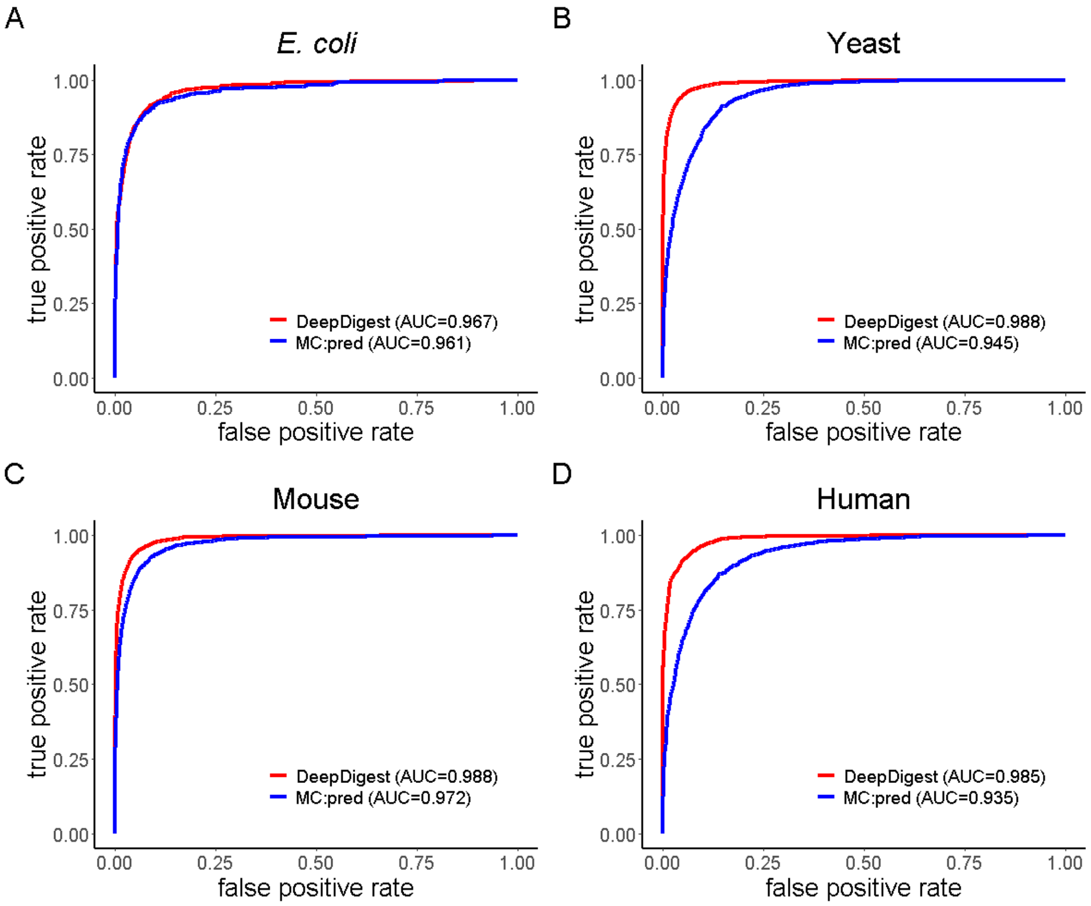


**Figure S1.** Performance comparison of DeepDigest and MC:pred on the four tryptic test sets from four different organisms (A) *E. coli*, (B) Yeast, (C) Mouse and (D) Human.

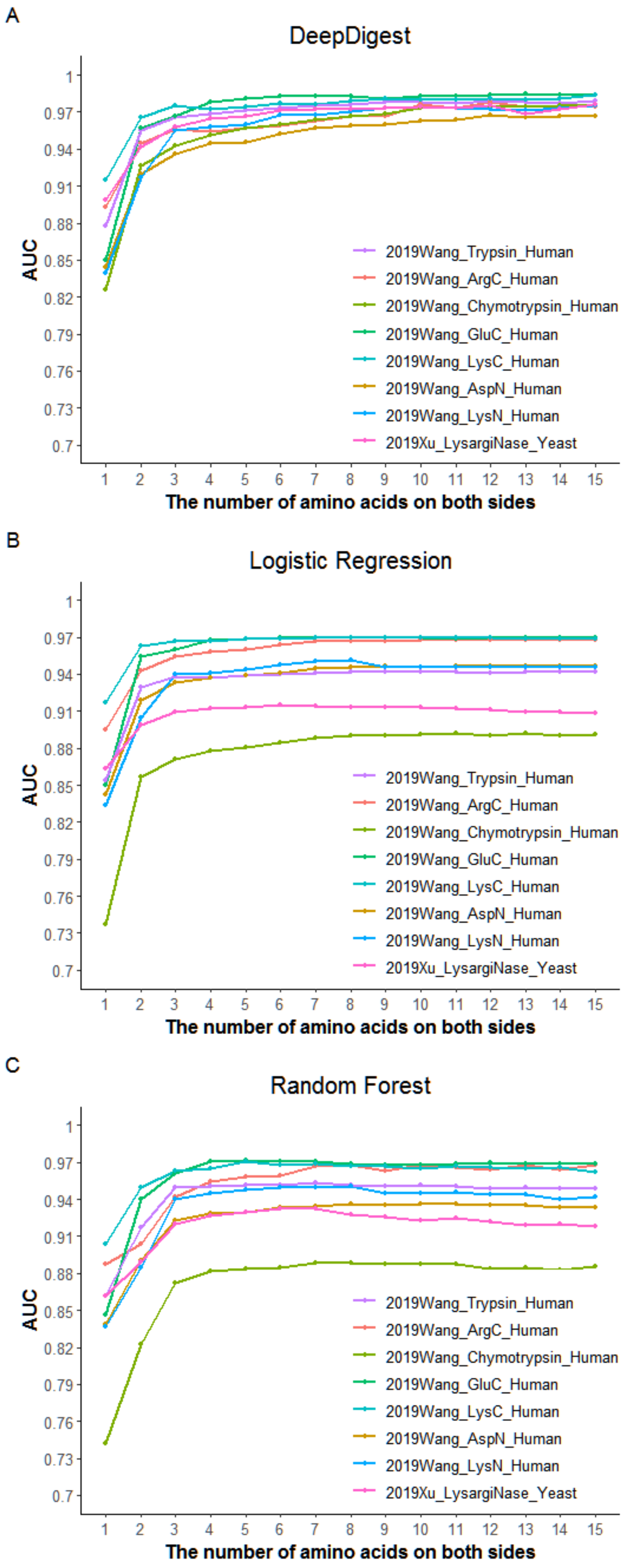


**Figure S2.** The AUCs of (A) DeepDigest, (B) Logistic Regression (LR) and (C) Random Forest (RF) with the increasing number of amino acids from both sides of the cleavage sites on the random 80-20% train-test splitting training sets.

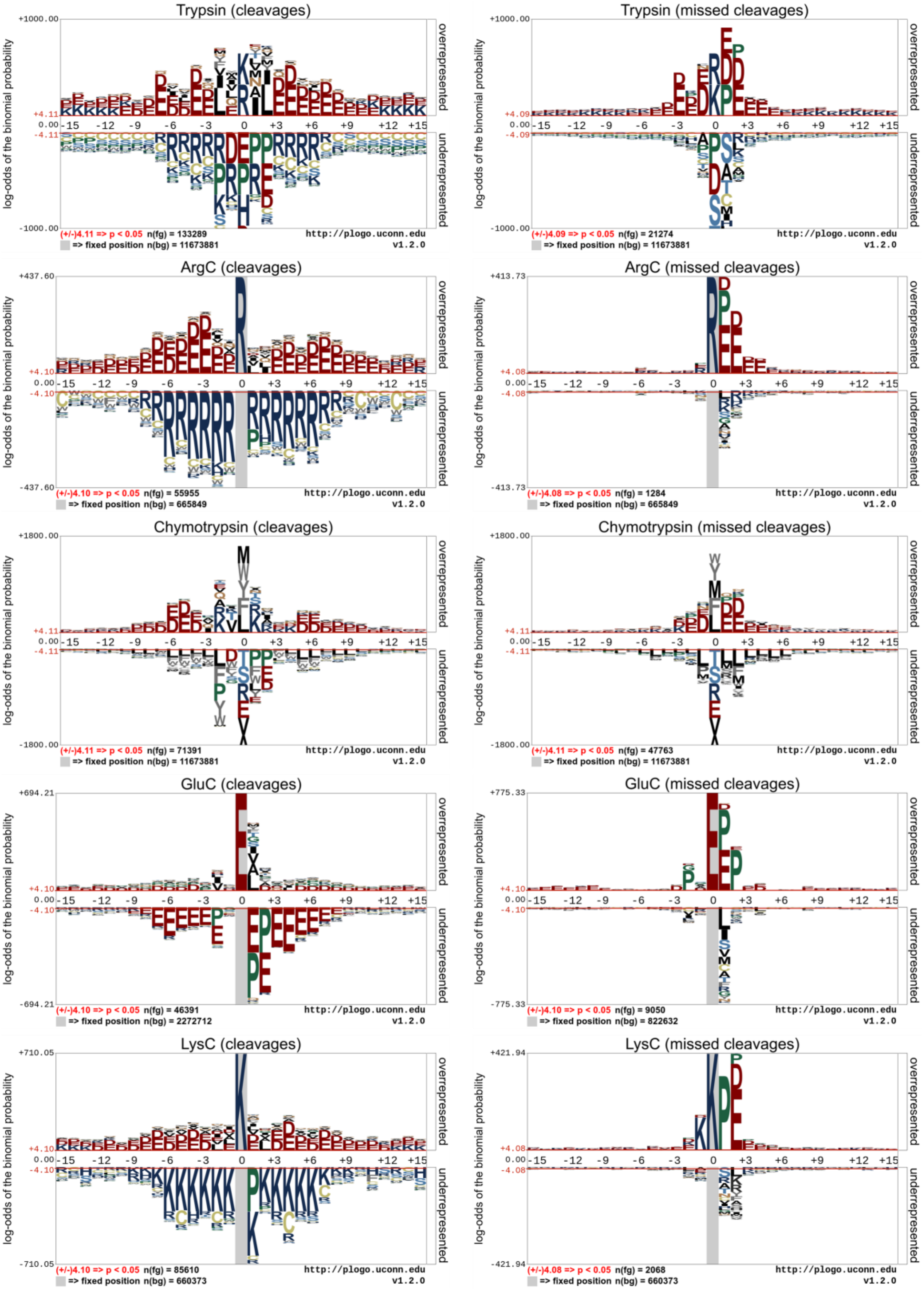
Trypsin (cleavages)
+1000.00
-1000.00
(+/-)4.11 => p < 0.05 n(fg) = 133289
=> fixed position n(bg) = 11673881
http://plogo.uconn.edu
v1.2.0
Trypsin (missed cleavages)
+1000.00
-1000.00
(+/-)4.09 => p < 0.05 n(fg) = 21274
=> fixed position n(bg) = 11673881
http://plogo.uconn.edu
v1.2.0
ArgC (cleavages)
+437.60
-437.60
(+/-)4.10 => p < 0.05 n(fg) = 55955
=> fixed position n(bg) = 665849
http://plogo.uconn.edu
v1.2.0
ArgC (missed cleavages)
+413.73
-413.73
(+/-)4.08 => p < 0.05 n(fg) = 1284
=> fixed position n(bg) = 665849
http://plogo.uconn.edu
v1.2.0
Chymotrypsin (cleavages)
+1800.00
-1800.00
(+/-)4.11 => p < 0.05 n(fg) = 71391
=> fixed position n(bg) = 11673881
http://plogo.uconn.edu
v1.2.0
Chymotrypsin (missed cleavages)
+1800.00
-1800.00
(+/-)4.11 => p < 0.05 n(fg) = 47763
=> fixed position n(bg) = 11673881
http://plogo.uconn.edu
v1.2.0
GluC (cleavages)
+694.21
-694.21
(+/-)4.10 => p < 0.05 n(fg) = 46391
=> fixed position n(bg) = 2272712
http://plogo.uconn.edu
v1.2.0
GluC (missed cleavages)
+775.33
-775.33
(+/-)4.10 => p < 0.05 n(fg) = 9050
=> fixed position n(bg) = 822632
http://plogo.uconn.edu
v1.2.0
LysC (cleavages)
+710.05
-710.05
(+/-)4.10 => p < 0.05 n(fg) = 85610
=> fixed position n(bg) = 660373
http://plogo.uconn.edu
v1.2.0
LysC (missed cleavages)
+421.94
-421.94
(+/-)4.08 => p < 0.05 n(fg) = 2068
=> fixed position n(bg) = 660373
http://plogo.uconn.edu
v1.2.0
log-odds of the binomial probability
overrepresented
underrepresented
-15 -12 -9 -6 -3 0 +3 +6 +9 +12 +15

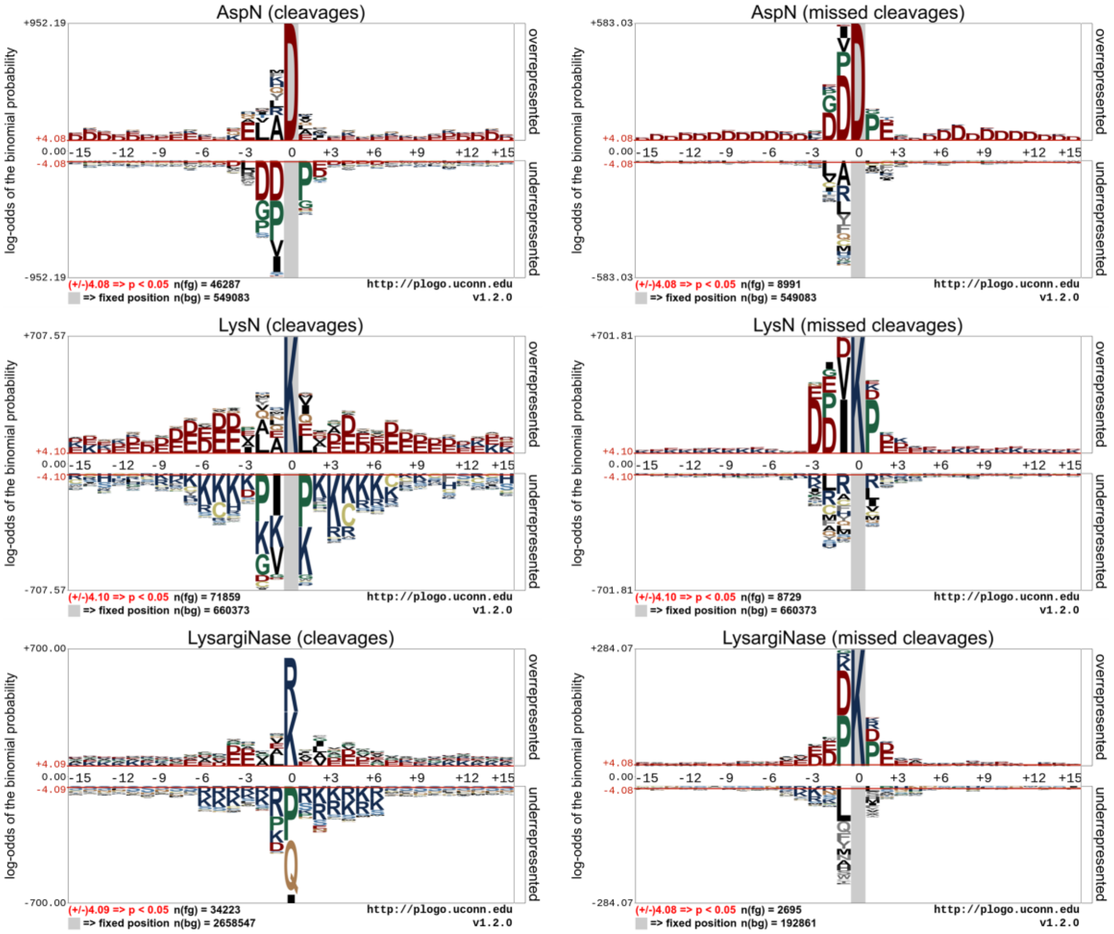


**Figure S3.** Sequence logos generated by pLogo. The red horizontal lines denote the threshold of $p<0.05$. Residues in a pLogo motif figure are scaled proportional to their log-odds binomial probabilities.

### Supplementary Tables

**Table S1.** The representation of each amino acid.

| **Amino acid** | **1-Letter** | **Code** |
|---|---|---|
| Alanine | A | 1 |
| Cysteine | C | 2 |
| Aspartic | D | 3 |
| Glutamic | E | 4 |
| Phenylalanine | F | 5 |
| Glycine | G | 6 |
| Histidine | H | 7 |
| Isoleucine | I | 8 |
| Lysine | K | 9 |
| Leucine | L | 10 |
| Methionine | M | 11 |
| Asparagine | N | 12 |
| Proline | P | 13 |
| Glutamine | Q | 14 |
| Arginine | R | 15 |
| Serine | S | 16 |
| Threonine | T | 17 |
| Valine | V | 18 |
| Tryptophan | W | 19 |
| Tyrosine | Y | 20 |
| Illegal Amino Acids | X, B, U, J, Z, O | 21-26 |

**Table S2.** Ten-fold cross-validation results of DeepDigest on the eight training sets using three metrics (AUC, F1 score and MCC), compared with logistic regression (LR), random forest (RF), and support vector machine (SVM).

| Dataset | Model | AUC | F1 Score | MCC |
|---|---|---|---|---|
| 2019Wang_Trypsin_Human | **DL** | **0.980** | **0.844** | **0.820** |
| | LR | 0.943 | 0.708 | 0.677 |
| | RF | 0.951 | 0.621 | 0.628 |
| | SVM | 0.949 | 0.741 | 0.712 |
| 2019Wang_ArgC_Human | **DL** | **0.970** | **0.639** | **0.631** |
| | LR | 0.965 | 0.576 | 0.574 |
| | RF | 0.960 | 0.013 | 0.079 |
| | SVM | 0.965 | 0.572 | 0.575 |
| 2019Wang_Chymotrypsin_Human | **DL** | **0.976** | **0.904** | **0.839** |
| | LR | 0.890 | 0.759 | 0.610 |
| | RF | 0.886 | 0.729 | 0.589 |
| | SVM | 0.898 | 0.768 | 0.628 |
| 2019Wang_GluC_Human | **DL** | **0.983** | **0.860** | **0.835** |
| | LR | 0.967 | 0.806 | 0.771 |
| | RF | 0.967 | 0.765 | 0.737 |
| | SVM | 0.969 | 0.794 | 0.759 |
| 2019Wang_LysC_Human | **DL** | **0.977** | **0.736** | **0.730** |
| | LR | 0.966 | 0.643 | 0.642 |
| | RF | 0.962 | 0.284 | 0.392 |
| | SVM | 0.959 | 0.697 | 0.697 |
| 2019Wang_AspN_Human | **DL** | **0.968** | **0.805** | **0.766** |
| | LR | 0.950 | 0.729 | 0.682 |
| | RF | 0.938 | 0.571 | 0.564 |
| | SVM | 0.953 | 0.744 | 0.699 |
| 2019Wang_LysN_Human | **DL** | **0.974** | **0.799** | **0.775** |
| | LR | 0.951 | 0.717 | 0.686 |
| | RF | 0.945 | 0.423 | 0.478 |
| | SVM | 0.952 | 0.722 | 0.694 |
| 2019Xu_LysargiNase_Yeast | **DL** | **0.971** | **0.719** | **0.694** |
| | LR | 0.917 | 0.497 | 0.468 |
| | RF | 0.923 | 0.079 | 0.185 |
| | SVM | 0.914 | 0.488 | 0.471 |

**Table S3.** Test results of DeepDigest on the eleven independent test sets using three metrics (AUC, F1 score and MCC), compared with logistic regression (LR), random forest (RF), and support vector machine (SVM).

| **Training Data** | **Test Data** | **Model** | **AUC** | **F1 Score** | **MCC** |
|---|---|---|---|---|---|
| 2019Wang_Trypsin_Human | 2016Schmidt_Trypsin_Ecoli | DL | **0.949** | 0.593 | 0.589 |
| | | LR | 0.916 | 0.552 | 0.539 |
| | | RF | 0.924 | **0.620** | **0.601** |
| | | SVM | 0.919 | 0.554 | 0.544 |
| | 2014Hebert_Trypsin_Yeast | **DL** | **0.975** | **0.822** | **0.796** |
| | | LR | 0.936 | 0.696 | 0.662 |
| | | RF | 0.943 | 0.616 | 0.624 |
| | | SVM | 0.940 | 0.718 | 0.685 |
| | 2016Malmström_Trypsin_Mouse | **DL** | **0.977** | **0.724** | **0.712** |
| | | LR | 0.947 | 0.666 | 0.642 |
| | | RF | 0.961 | 0.704 | 0.680 |
| | | SVM | 0.956 | 0.684 | 0.664 |
| | 2019Miller_Trypsin_Human | **DL** | **0.974** | **0.806** | **0.778** |
| | | LR | 0.929 | 0.671 | 0.633 |
| | | RF | 0.958 | 0.720 | 0.697 |
| | | SVM | 0.946 | 0.741 | 0.708 |
| 2019Wang_ArgC_Human | 2019Miller_ArgC_Human | **DL** | **0.960** | **0.497** | **0.510** |
| | | LR | 0.944 | 0.475 | 0.481 |
| | | RF | 0.956 | 0.378 | 0.374 |
| | | SVM | 0.946 | 0.475 | 0.481 |
| 2019Wang_Chymotrypsin_Human | 2019Miller_Chymotrypsin_Human | **DL** | **0.969** | **0.872** | **0.793** |
| | | LR | 0.878 | 0.737 | 0.583 |
| | | RF | 0.937 | 0.811 | 0.705 |
| | | SVM | 0.921 | 0.802 | 0.687 |
| 2019Wang_GluC_Human | 2019Miller_GluC_Human | **DL** | **0.958** | **0.773** | **0.730** |
| | | LR | 0.946 | 0.678 | 0.646 |
| | | RF | 0.936 | 0.661 | 0.637 |
| | | SVM | 0.951 | 0.692 | 0.663 |
| 2019Wang_LysC_Human | 2019Miller_LysC_Human | DL | 0.842 | **0.382** | **0.392** |
| | | LR | **0.851** | 0.306 | 0.331 |
| | | RF | 0.805 | 0.239 | 0.310 |
| | | SVM | 0.835 | 0.336 | 0.360 |
| 2019Wang_AspN_Human | 2019Miller_AspN_Human | DL | 0.963 | 0.563 | 0.570 |
| | | LR | 0.965 | 0.559 | 0.566 |
| | | RF | 0.962 | **0.632** | **0.619** |
| | | SVM | **0.972** | 0.572 | 0.582 |
| 2019Wang_LysN_Human | 2018Zhang_LysN_Ecoli | DL | **0.809** | 0.341 | 0.289 |
| | | LR | 0.736 | 0.335 | 0.267 |
| | | RF | 0.779 | **0.421** | **0.398** |

| | | | | | |
|---|---|---|---|---|---|
| | | SVM | 0.744 | 0.337 | 0.269 |
| 2019Xu_LysargiNase_Yeast | 2018Zhang_LysargiNase_Ecoli | **DL** | **0.872** | **0.454** | **0.428** |
| | | LR | 0.770 | 0.336 | 0.317 |
| | | RF | 0.803 | 0.111 | 0.218 |
| | | SVM | 0.788 | 0.311 | 0.315 |
